# SNNs Are Not Transformers (Yet): The Architectural Problems for SNNs in Modeling Long-Range Dependencies

**DOI:** 10.1101/2025.10.31.685901

**Authors:** William Fishell, Suraj Honnuraiah

## Abstract

Spiking neural networks (SNNs) have attracted growing interest for their ability to operate efficiently on low-power neuromorphic hardware, offering a biologically grounded route toward energy-efficient computation. However, despite advances in large-scale neuromorphic systems capable of simulating millions of spiking neurons and synapses, SNNs continue to underperform state-of-the-art (SOTA) artificial neural networks (ANNs) on complex sequence-processing tasks.

Here, we present an explicit covering-number bound analysis for SNNs based on the non-leaky integrate and fire (nLIF) model. Leveraging recent work on causal partitions and local Lipschitz continuity, we derive a global Lipschitz constant and show that the sample complexity of nLIF networks scales quadratically with input sequence length. We analytically compare these bounds with those of Transformer and recurrent neural network (RNN) architectures, revealing fundamental constraints on how current SNNs process long-range dependencies. Finally, we show that these theoretical assumptions align with known cortical mechanisms, particularly inhibitory normalization and refractoriness, and discuss their implications for developing future neuromorphic architectures that more closely approximate biological computation.

## Introduction

References to biology have been important in pushing the bounds of machine learning models. From the application of the Hebbian learning rule in Hopfield networks to the convolutional filters that emulate local receptive fields in the visual cortex, modern deep-learning models continually draw practical inspiration from biological systems. Although strict biological realism isn’t required—many innovations like self-attention and gradient descent have no direct counterpart in real neurons—grounding our algorithms in biological principles offers clear benefits: by drawing on decades of neuroscience research, we can uncover efficient mechanisms for learning and adaptation, and by emulating evolution’s optimizations for energy use, robustness, and adaptability, we can design models that are both powerful and resource-efficient.

Spiking Neural Networks (SNN)—often called the “third generation” of neural models—offer greater biological realism than standard artificial neural networks (ANN). These networks achieve remarkable energy efficiency by emitting sparse, temporally distributed spikes—much like biological action potentials. Their built-in sparsity and sequential processing stand in stark contrast to the *O*(*N* ^2^) complexity of Transformer self-attention and the billions of operations required during large-language-model inference. SNNs have already achieved promising results, such as SpikeGPT, which achieves performance comparable to GPT-3 while using 33 times less power [1]. Furthermore, these networks leverage both temporal and spatial dependencies and include a richer set of programmable parameters—such as delays and neuron-specific thresholds—which endows them with greater expressiveness than recurrent neural networks of the same depth and width [2, 3].

Despite the potential for increased expressivity, SNNs struggle from many of the same limitations that RNNs have: a finite memory horizon [4], and the inability to model non-sequential interaction [5]. These issues in SNNs have caused them to struggle with modeling long-term dependencies. Creating wider adoption of these networks requires devising modifications that solve these issues. Inspired by Edelman et al. [6], we use a learning-theoretic approach to understand how learning varies with sequence length. Measuring the change in the number of samples required to achieve PAC learnability as a function of the sequence length of those samples indicates how well the SNN architecture models long-range dependencies. While prior work on the VC-dimension of SNNs has established foundational capacity bounds [7, 2, 8, 9], it has largely overlooked the role of sequence length and has not explored covering-number approaches. In this work, we leverage recent results on the local Lipschitz continuity of SNNs [10] to derive a covering-number bound, establishing a novel relationship between sample complexity and sequence length. Our analysis provides new theoretical insights into SNNs and helps explain their current limitations in language modeling tasks compared to Transformers and RNNs.

An SNN processes information as time-stamped spike trains instead of static real-valued activations. Each neuron integrates incoming weighted spikes from the preceding layer, accumulating a membrane potential: this potential decays continuously. When it exceeds a fixed threshold, the neuron emits a spike and transmits its associated edge weights to downstream connections after a time delay Δ*t*. This characteristic of the SNN results in sparse network activity. As each spike train passes through, only some of the neurons fire. Because the computation unfolds in continuous time, the network’s state —and thus its output —depend on the moment of observation by the output layer. We focus on the non-leaky and leaky integrate-and-fire (nLIF, LIF) neuron models—the most widely adopted spiking neuron models. Although all SNNs share the pipeline sketched above, they diverge in their spike-encoding schemes and in the synaptic delays they assume. The LIF model captures these ingredients in their simplest canonical form and underlies many of the more sophisticated spiking architectures encountered in practice (for a more detailed overview of the specifics of SNNs and LIFs, see App. A). The nLIF is a simpler form of the LIF that does not have a leaky membrane potential (for a formal definition of nLIFs see App. C).

## Results

### 1 Pseudo-Attention in the Leaky Integrate-&-Fire Model

The LIF architecture enables earlier tokens to influence later ones by effectively ‘attending’ to them and thus modifying the state of the network^1^. Although this mechanism is not identical to traditional attention, several similarities are noteworthy. At its simplest, an attention mechanism quantifies the similarity between pairs of tokens in the context. Suppose tokens *c*_1_ and *c*_2_ occur at positions *i* and *i* + *n*, so

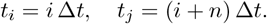

We represent their embeddings as instantaneous spike-train vectors.

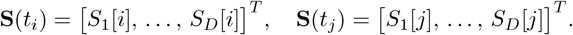

The scaled dot product does not occur in the LIF, but at **S**(*t*_*i*_), the spike train raises the membrane potentials of its active subset of neurons, so when **S**(*t*_*j*_) arrives, the neurons that represent these two tokens are more likely to fire. In contrast, if the overlap is small, fewer neurons will spike at time *t*_*j*_ representing their similarity. The state of the network at any time *t* is defined as a function of both the temporal dynamics and the contextual input, where the interactions between all pairs *c*_*i*_ and *c*_*j*_ are inherently present. Based on the described behavior, the LIF architecture strikes a balance between an RNN and a Transformer in expressing non-sequential interactions. Unlike an RNN, the LIF model supports some non-sequential computation. However, its inherent sequential nature — stemming from how tokens feed into the network as spike trains — makes it impossible to realize the attention mechanism fully. Furthermore, the introduction of distance between two tokens can introduce noise, which can affect the network state and obscure comparisons between tokens in the LIF architecture, thereby acting as a further barrier to attention in SNNs.

Conceptually, the LIF architecture can be viewed as having a kind of short-term memory. Because each neuron’s membrane potential carries over across time steps, it retains a faint trace of recent activity. That carry-over creates limited non-sequential interactions, enough for nearby events to influence one another, though without forming true recurrence. In this sense, LIF neurons offer a weak analog of attention, providing selective temporal weighting without explicit dot-product computations, in contrast to the query–key matching used in Transformers. This produces short-range contextual effects that resemble attention kernels in their temporal weighting but remain strictly sequential in operation. Variability in spike timing with temporal distance can introduce noise that may obscure these effects, and a quantitative comparison with attention mechanisms is left for future work.

### 2 Sample Complexity of SNNs, Transformers, and RNNs

The VC dimension of a feedforward spiking neural network with *K* synaptic edges grows tightly on the order of Θ(*K*^2^) [11, 8]. Intuitively, this quadratic scaling reflects the vast combinatorial flexibility afforded by programmable parameters—each weight, delay, and threshold setting contributes to carving out distinct decision regions, allowing the network to shatter on the order of *K*^2^ input points. Although Maass’ original proof focused on the nLIF model, the same argument holds for the standard LIF architecture: introducing additional programmable leak parameters can only maintain or increase the network’s expressive capacity. Furthermore, this does not increase the VC dimension of the SNN, as the leaks function simply as another programmable parameter (for more details on why this is the case, see App. B). There is a strong understanding of the expressive capacity of the SNN, yet comparatively little research on how this capacity changes as the size of the inputs being modeled changes. Transformers and RNNs have well-understood bounds on how the number of samples required to achieve PAC learnability scales with the input sample size. RNNs require linear growth in the size of an input in the number of samples to achieve PAC learnability as the sequence size grows. Transformers achieve an even more impressive *log*(*T*) sample increase requirement for modeling longer sequences [6, 12]^1^. Unlike Transformers, which have an inductive bias towards sparsity, SNNs have an inductive bias towards low-frequency functions [13]. This inductive bias indicates that they may perform poorly on modeling the relationship between a small set of important input tokens: the long-range dependency modeling task. To date, no analysis has quantified how SNN sample complexity grows with sequence length. In this work, we fill that gap by deriving explicit bounds for nLIF networks, revealing exactly how samples must scale as sequences get longer. These insights deepen our theoretical understanding of spiking architectures and help explain their inherent bias toward low-frequency mappings.

#### 2.1 Non Leaky Integrate-&-Fire Covering Number

Under appropriate assumptions on our function class, we show that an nLIF SNN is globally Lipschitz continuous. We then derive the covering-number for both single-token and multi-token inputs and compare them to quantify how sample complexity scales with sequence length. Henceforth, we refer to the single-input-token setting as the Single Input Spike (SIS) problem and the multi-input-token setting as the Multi Input Spike (MIS) problem.

#### 2.2 SIS Covering Number

The output spike time of an nLIF neuron can be made locally Lipschitz continuous by restricting attention to the specific subset of its inputs and weights that actually cause it to fire. From this, we show, with proper assumptions, the global Lipschitz constant and derive the covering number for the SIS case, and then extend to the MIS problem. Our work builds on work that proves local Lipschitz continuity with respect to weight and input spike time for an output spike time in an nLIF SNN. In deriving the covering number, we introduce common terminology used for the remainder of the paper.

A **causal piece**, denoted as P, for neuron *i* in layer *ℓ* is the set of all pre-synaptic spike-time vectors and weights that lead to its firing:

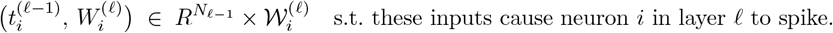

Its **causal set** is the set of pre-synaptic neuron spike times whose spikes precede its firing time:

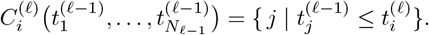

Lastly a **causal path**, denoted 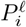, for neuron *i* in layer *ℓ* is the set

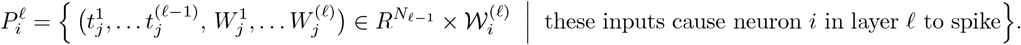

Recent work shows that, when restricted to a fixed causal piece, the output spike-time map is locally Lipschitz continuous in those pre-synaptic times and weights^2^ [10] (For a proof sketch and further details on this paper, see Appendix C).

Given a causal set for some neuron in layer *ℓ* that spikes at time *T*, the causal set is locally Lipschitz continuous with respect to its causal piece.

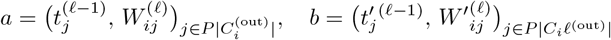

be two configurations of pre-synaptic spike times and weights on the causal piece 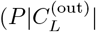 and satisfy

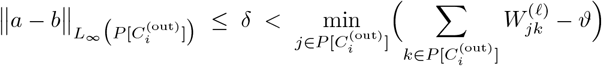

So that any perturbation of size at most δ neither delays an originally–causal pre-synaptic spike past the output time *T* nor advances a non-causal spike before *T*. The total weighted input stays above the threshold, so the neuron still fires at *T*. Then the output spike-time map 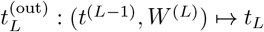 is locally Lipschitz continuous on that piece, with

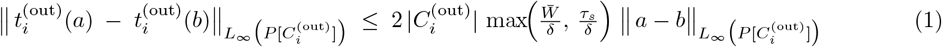

where

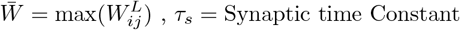

##### Assumption 2.1

(Assumptions). *Let*

1. *the network be a feedforward nLIF SNN;*
2. *M be a set of training examples s*.*t*.

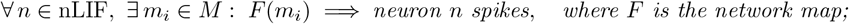
3. *each neuron’s causal set 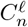 is non-empty;*
4. *all synaptic weights 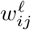 are positive and, for every causal piece P*,

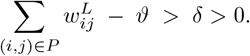
5. *the reset for a neuron’s membrane after spiking is instantaneous*

##### Theorem 2.1

(Global Lipschitz Bound). *Under Assumption 2*.*1, there exists a constant L*_global_ *s*.*t. for any two input-spike patterns a, b with* 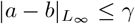.

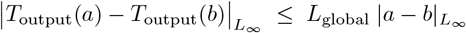

*Proof*. Consider the output layer of neurons in the nLIF. By the assumption, each of the neurons in the output layer has some causal set 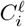 that is not empty. Because ∃ a δ margin across all the thresholds and the causal set of each of these output neurons is non-empty, we can construct a Layer Lipschitz bound across this layer by taking the

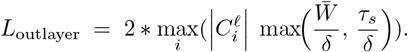

thus any pre-synaptic inputs a,b 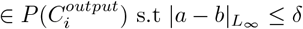 are locally Lipschitz continuous for neuron 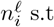.

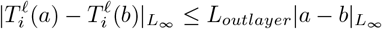

This can be traced recursively through the layers towards the input by telescoping. Consider *a, b* are now input spikes in the causal piece of neuron 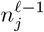 and *a, b* 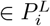 s.t

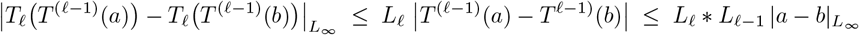

with the new bounds on the input that 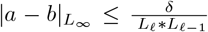 where *L*_*ℓ*_ and *L*_*ℓ*−1_ are defined as the max Lipschitz constants of their respective layers. By continuing this process to the input layer, we can construct a global Lipschitz constant for the entire nLIF feedforward network.

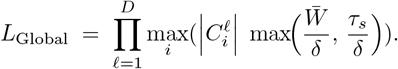

Because any input spikes *a, b* are in some causal path and all neurons in the network are covered in some causal path s.t. *L*_*global*_ can bound the output spike times of that neuron, *L*_*global*_ is a global Lipschitz constant with respect to spike time and weight for the network. By simply setting our free term 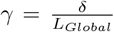, the differences in the inputs will not cause a change in any causal path, so the entire network is defined by the global Lipschitz constant *L*_*global*_

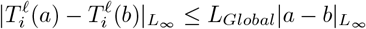

Under our assumptions, the nLIF network is globally Lipschitz continuous in the spike-time domain. To extend this to real-valued inputs and outputs, we compose three Lipschitz maps: an encoder that converts each real input into a spike train, the nLIF network itself (which is Lipschitz in spike times), and a decoder that reads the output spike train and maps it back to the target space. Because the composition of Lipschitz maps remains Lipschitz, the end-to-end map from original inputs to final outputs is Lipschitz. This lets us apply standard covering-number bounds directly in the real-valued signal domain. To illustrate the simplicity of constructing such a composition, we use a toy example of classifying words (for the details of this Example see App. D).

**Example D** outlines a process of applying a time to first spike (TTFS) encoding and softmax decoding to the input and output spaces to create a global Lipschitz constant for the function. Using the composition of Lipschitz continuous functions and the results from **Theorem ??**, a function class *F* can be created from nLIF SNN, and a covering number can be constructed. Provided the assumptions in 2.1 are met. For each of these functions in the function class, there is a global Lipschitz constant defined as

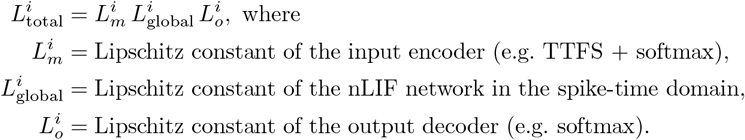

By taking the *max* 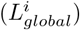, a Lipschitz constant which covers the entire function class can be derived, denoted *L*_*MaxT otal*_. Due to our restriction on distinguishing perturbations of at most *γ*, the covering number is

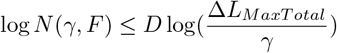

This bound enables us to control the error within the function, and thus, the sample complexity M becomes

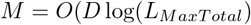

Crucially, *L*_MaxTotal_ scales with each layer’s maximum causal-set size. Allowing neurons to fire multiple times (MIS) rather than just once (SIS) thus induces a pronounced jump in the resulting sample complexity, because the causal sets grow with respect to the number of input spikes.

#### 2.3 MIS Covering Number

The MIS problem consists of a sequence of tokens inputted as a spike train, where each token follows the preceding element after some time delay. Building on **Example D**, we encode a sequence of tokens into spike times as follows:

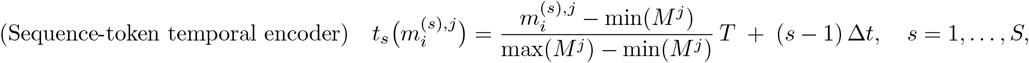

This encoding scheme can be implemented using a Dirac delta function to fire pulses at the moments of interest, as encoded by the sequence-token temporal encoder.

We follow the SIS setup described in **Example D** by keeping *T* small enough that a single token can trigger at most one spike per neuron as it moves through the network. Over a sequence of tokens, however, a neuron may fire multiple times.

##### Theorem 2.2

(Global Lipschitz Bound for MIS). *Under Assumption 2*.*1, there exists a constant L*_globalS_ *s*.*t. for any two multi-spike input patterns a, b of length S*,

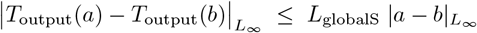

*Here, L*_globalS_ *is the global Lipschitz constant for the sequence-based encoding of length* |*S*|, *and it satisfies*

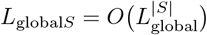

*Where L*_global_ *is the global Lipschitz constant established for the SIS problem on the same nLIF SNN*.

This result can be extended by using a composition of Lipschitz-continuous functions to encode and decode the spike train, as in **Example D**. In particular, for any sequence *S* ∈ *M* ^*S*^, applying the nLIF under suitable input bounds yields a globally Lipschitz map

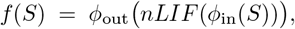

with constant *L*_*Func*_ s.t.

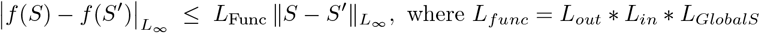

##### Theorem 2.3.

*For any nLIF network for which we can establish global Lipschitz continuity, the worst-case sample complexity grows quadratically with the sequence length*.

*Proof*. Given a function class *F*) For the MIS problem, a global Lipschitz constant can be constructed for each of the *f*_*i*_ in the function class. By taking the maximum of these Lipschitz constants and invoking

**Theorem 2.2**, the maximum global Lipschitz constant is equal to some 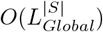 where *L*_*Global*_ is the constant for the SIS case. Thus, the covering number is

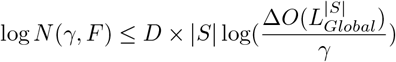

When |*S*| is the sequence length, applying standard log rules, |*S*| can be brought down. Thus, the sample complexity for this problem is

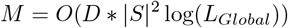

Controlling for the input dimension *D* (the size of each tensor) and the network’s structure by increasing the sequence length, we need |*S*|^2^ more samples to ensure PAC learnability.

In an nLIF SNN, the *D* input channels arrive as an *S*-length spike train whose contributions aggregate into each neuron’s membrane. In the MIS setting, the global Lipschitz constant compounds by a factor of *L*_SIS_ at each of the *S* spikes—growing as

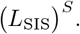

Since the covering-number bound also scales linearly with the *DS*-dimensional input space, this multiplicative Lipschitz blow-up combined with the dimension term yields the sample-complexity penalty.

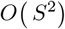

The quadratic sample-complexity growth both illustrates why nLIF SNNs struggle with long-term dependencies and shows that preventing explosive causal set expansion can improve their ability to model such dependencies. By preventing unbounded excitation, we can cap causal set expansion, thereby bounding quadratic sample-complexity growth.

Causal pieces are an active area of research, and it is currently not known whether causal pieces in SIS or MIS LIFs are locally Lipschitz continuous with respect to their causal piece. We leave the question of sample complexity for an LIF to a later paper, as further research is needed to understand how the causal set of a neuron grows when it is leaking.

### 3 Implications for Biological Neural Circuitry

The spiking dynamics of nLIF SNNs lead to weak theoretical guarantees for learning long-range dependencies. By mapping the architectural assumptions of these networks to their biological counterparts, we both demonstrate the biological plausibility of our framework and identify which additional neural mechanisms may be required in real circuits to achieve stronger learning guarantees.

#### 3.1 Effect of Refractory Gating and Population Normalization

The refractory period following Na^+^ activation prevents uncontrolled excitation in neural circuits ([14], [15]). Despite endogenous limits on neuronal firing, the sequential integration of inputs leads to quadratic scaling.

##### Lemma 3.1

(Refractory Period Constraint). *Let τ*_*ref*_ > 0 *be the refractory period enforcing* Δ*t*_min_ ≥ *τ*_*ref*_ *between consecutive spikes of neuron i. For a sequence of length S with inter-token delay* Δ*t, the maximum spike count is:*

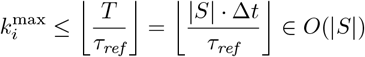

*Despite this constraint, the global Lipschitz constant satisfies:*

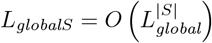

*and therefore sample complexity remains:*

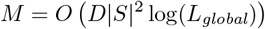

*Proof*. A neuron can fire at most once per *τ*_ref_ interval. Over total time *T* = |*S*| · Δ*t*, this bounds *k*_max_ ∈ *O*(|*S*|). However, in the MIS proof (Theorem 2.2), causal sets grow as 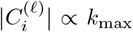 across the sequence. Thus:

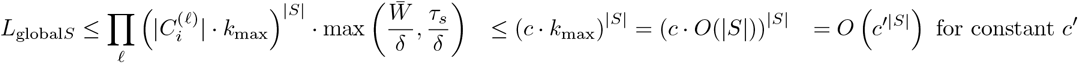

The exponential structure is preserved.

Lemma 3.1 demonstrates that the refractory period does not induce inactivity in a manner that gets rid of the exponential |*S*|. Therefore, quadratic scaling is preserved.

##### Lemma 3.2

(Population Normalization - Gain Control). 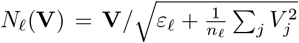 *be a divisive normalization operator with bounded Jacobian* 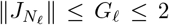. *Normalization ([16], [17]) modifies per-layer bounds by a constant factor:*

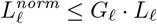

*L*_*ℓ*_ *but preserves the exponential structure for sequences:*

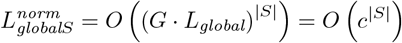

*Therefore, sample complexity remains quadratic: M* = *O*(*D* |*S* |^2^ log(*L*_*global*_)). *This aligns with cortical divisive normalization, demonstrating that biological gain control reduces constants but does not linearize scaling*.

At first glance, this behavior appears to be a network saturation problem — something biological systems would avoid through population-level normalization. However, as shown in Lemma 3.2, the sequential manner in which these signals are integrated still preserves the exponential growth structure and quadratic scaling, despite normalization.

*Proof of Lemma 3*.*2*. The divisive normalization operator *N*_*ℓ*_ has bounded Jacobian 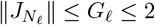. For a normalized layer 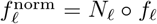, the chain rule gives 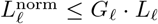. Following Theorem 2.2 telescoping argument for sequence composition:

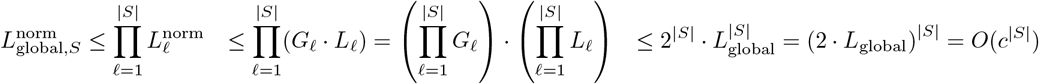

By Theorem 2.3, *M* = *O*(*D* |*S*|^2^ log(*c*^|*S*|^)) = *O*(*D* |*S*|^2^) since log(*c*^|*S*|^) = |*S*| log *c* absorbs into the quadratic term. The exponential structure is preserved, demonstrating that biological gain control reduces constants but does not linearize scaling.

Because the brain is remarkably efficient at modeling language and learning long-range dependencies, we hypothesize that a third mechanism is present in biological neural nets that enables efficient learning by bounding uncontrolled excitation.

#### 3.2 Necessity of Inhibitory Structures

Lateral inhibition linearizes the quadratic scaling induced by spiking dynamics, enabling the brain to efficiently learn long-range dependencies.

##### Lemma 3.3

(Inhibitory Sparsification and Telescoping MIS Bound). *Let* 𝒩 *be a network with L layers processing sequences of length* |*S*|. *Consider two inhibitory mechanisms ([18]):*

1. ***Per-bin WTA cap:*** *A hard limit s*_*ℓ*_ *on the number of active neurons per layer per time bin*.
2. ***Threshold boost:*** *Dynamic threshold adjustment*

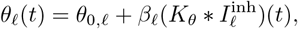

*where* 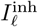 *represents inhibitory input (modeling PV/SST interneurons). Then the following hold:*

i. ***Margin bounds:***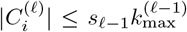 *where* 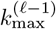 *is the maximum number of spikes per neuron in layer ℓ* − 1.
ii. ***Effective margin preservation:*** *The effective margin* 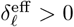 *remains bounded across layers*.
iii. ***Telescoping global Lipschitz bound:*** 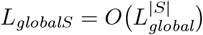*where L*_*global*_ *is the single-spike Lipschitz constant*.
iv. ***Sample complexity:*** *M* = *O D*|*S*|^2^ log *L*_*global)*_.

***Critical distinction:*** *Soft inhibitory mechanisms (WTA caps with s*_*ℓ*_ = Ω(|*S*|) *or threshold boosts) reduce only the* constants. *To achieve* linear *scaling, a hard constraint k*_max_ ≤ *K* = *O*(1) *independent of* |*S*| *is required*.

##### Lemma 3.4

(Linearization via H ard Per-Sequen ce Cap). *Impose a* ***hard constraint*** *k*_max_ ≤ *K* = *O*(1) *independent of* |*S*|. *Then M* = *O (D*|*S*| log *L*_*global)*_

*Proof*. Consider a neuron 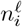 processing a sequence of |*S*| inputs. The first input *S*_1_ propagates through the resting network with local Lipschitz constant:

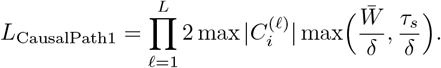

Subsequent inputs encounter neurons with residual membrane potential. The causal set for *S*_2_ depends on that of *S*_1_, causing interdependence:

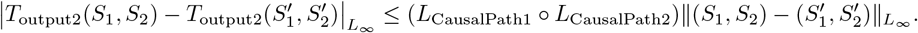

Without inhibition, causal sets accumulate contributions from all previous inputs. Telescoping through *S*_1_, …, *S*_|*S*|_ yields:

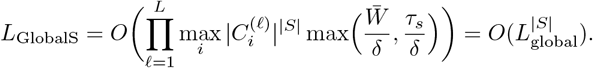

##### Hard cap breaks the telescoping

Impose *k*_max_ ≤ *K* = *O*(1): each neuron fires at most *K* times total across the entire sequence. For any subset of inputs {*S*_1_, …, *S*_*j*_} with *j* ≤ |*S*|:

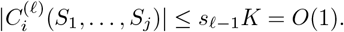

The total spike count is capped, so the per-layer Lipschitz factor remains *L*_layer,*ℓ*_ = *O*(1), and:

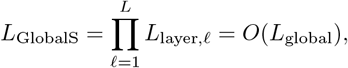

##### Sample complexity

The input diameter is diam(𝒳) = |*S*| · diam(𝒳_1_), but the effective dimension is *D*_eff_ = *O*(*D*_1_) due to the spike constraint:

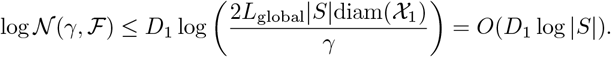

Taking 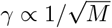 yields *M* = *O*(*D*_1_|*S*| log *L*_global_), linear in |*S*|.

To fit this biological explanation into our analysis of spiking dynamics, we must understand whether our derivation extends from nLIF networks to LIF networks. We are actively researching this extension, but hypothesize that the same behavior holds: we can always choose a Δ*t* for our rate encoding that is arbitrarily smaller than the leak time constant, effectively recovering nLIF-like behavior.

## Discussion

The quadratic scaling of sample complexity with sequence length in nLIF networks highlights their inherent difficulty in modeling long-term dependencies. Architectural factors — most notably the strictly sequential integration of inputs drives this growth; normalization can shrink constants but does not alter the sequence-length exponent. Importantly, our analysis shows that bounding the size of each neuron’s causal set is a direct route to improved sample efficiency as sequences lengthen. Future work should therefore explore implementations of inhibitory layers in SNNs. Lateral inhibition has shown promise in computer vision tasks [19], yet it remains to be tested in language modeling architectures.

Our analysis links common circuit phenomena to formal parameters that govern expressivity and efficiency in spiking networks. Inhibitory mechanisms, including effects consistent with PV- and SST-class interneuron activity, can reduce the size of active causal sets 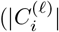; see Lemma 4.3), yielding sparser effective population codes. Divisive normalization–like gain control can bound per-layer sensitivity via factors *G*_*ℓ*_ (Lemma 4.2), thereby reducing Lipschitz constants *L*_*ℓ*_ without altering the sequence-length exponent. Refractory and leak dynamics (Lemma 4.1) cap the per-neuron spike count *k*_max_, limiting dynamic range but leaving the quadratic dependence on |*S*| unchanged unless an explicit hard cap is enforced.

These correspondences suggest that biologically plausible constraints help organize SNN dynamics within a Lipschitz-bounded regime that supports robustness and learnability. Together, these observations point toward the efficiency of biologically grounded, brain-inspired architectures, where inhibitory balance, normalization, and refractory dynamics jointly maintain robust and energy-efficient computation while preserving expressive capacity.

*Scope note*. All global Lipschitz and covering arguments presented here are proved for nLIF neurons. Extending to LIF will require establishing local Lipschitz continuity across reset surfaces, which we conjecture holds in the small-Δ*t* regime with bounded leak.

## Conclusion

We presented, to our knowledge, the first covering-number analysis of spiking architectures, establishing a quadratic sample-complexity bound for nLIF networks for monotone input sequences (MIS). This bound arises from the growth of causal sets across layers and time, which compounds global Lipschitz factors with sequence length. In this sense, sequential integrate-and-fire dynamics impose a structural limit on how spiking networks scale with long sequences.

Additionally, the analysis makes explicit how biologically plausible mechanisms map onto these formal parameters. Inhibitory interactions can sparsify active causal sets, divisive normalization can bound layer sensitivity and reduce Lipschitz constants, and refractory or leak dynamics can cap the per-neuron spike count. Together, these constraints shrink constants and, when a hard per-sequence inhibitory cap is enforced, can yield linear scaling in sequence length. This identifies concrete levers for architectural design and neuromorphic deployment.

While our results do not claim that spiking networks cannot capture long-range dependencies, they show that efficient sequence processing requires mechanisms that limit causal-set growth or otherwise regularize sensitivity. Assertions about frequency selectivity in language modeling remain hypotheses in this work and warrant targeted empirical tests.

All global Lipschitz and covering arguments here are proved for nLIF neurons. Extending these results to standard LIF models will require establishing local Lipschitz continuity across reset surfaces, which we conjecture holds in the small-Δ*t* regime with bounded leak.

More broadly, the correspondences developed here suggest that biologically grounded constraints help organize spiking network dynamics within a Lipschitz-bounded regime that supports robustness and learnability. This framework points toward brain-inspired, energy-efficient architectures in which inhibition, normalization, and refractory dynamics jointly maintain expressive yet stable computation.

## Author Contributions

W.F. conceived and performed the formal analysis, with S.H. contributing to conceptual design and biological contextualization. W.F. and S.H. jointly developed the methodological framework and discussed the theoretical implications. W.F. conducted the investigation, implemented analyses, and carried out mathematical derivations. S.H. provided supervision, critical feedback, and contextualized the biological relevance of the results. W.F. drafted the manuscript, and S.H. provided critical revision and editorial input. Both authors approved the final version of the manuscript.

## A Spiking Neural Networks

### Definition 1.

***Spiking Neural Networks (SNN)*** *are biologically plausible neural networks that evolve continuously in time and emit discrete spikes whenever the membrane potential exceeds a preset threshold. Their dynamics follow:*

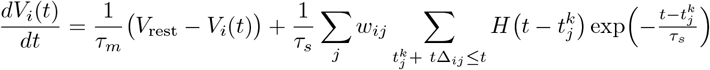

### Subject to the reset condition

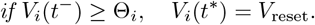

*τ*_*m*_ : Membrane time constant (leak rate).

*V*_*i*_(*t*) : Membrane potential of neuron *i* at time *t*.

*V*_rest_ : Resting (leak) potential.

*w*_*ij*_ : Synaptic weight from neuron *j* to neuron *i*.

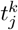 : Time of the *k*-th spike from neuron *j*.

*T*_*s*_ : Synaptic delay before post-synaptic effect.

*H*(·) : Heaviside step (spike) function.

*τ*_*s*_ : Synaptic time constant in *α*(*s*).

Θ_*i*_ : Firing threshold for neuron *i*.

*t*Δ_*ij*_ : The time it takes for a current to reach a postsynpatic neuron from a pre-synaptic neuron.

*V*_reset_ : Reset potential after a spike.

∑_*j*_ *w*_*ij*_ External input current to neuron *i* from all neurons in previous layer.

### A.1 Leaky Integrate & Fire Model

There are many biologically realistic neuronal models, but the Leaky Integrate & Fire Model (LIF) is the most commonly used due to its simplicity. The LIF simulates neuronal behavior by integrating incoming currents, applying a gradual leak over time, and emitting a spike when the membrane potential exceeds a set threshold. Unlike traditional artificial neural networks where neurons are updated synchronously, in the LIF model, neurons accumulate input over time and fire only when sufficient input is gathered to cross the threshold. The definition used for an SNN follows the same subthreshold dynamics of the membrane potential in an LIF model:

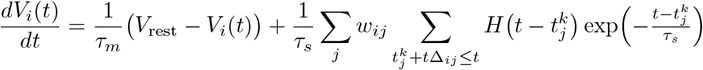

At time *t*, if the pre-synaptic neuron *j* at time 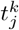 reaches it’s threshold, an instantaneous spike is triggered causing a jump in input current that then decays exponentially over the synaptic time constant *τ*_*s*_. Writing this out:

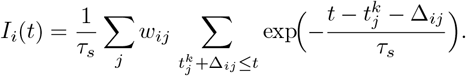

Each pre-synaptic spike at time 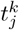 reaches neuron *i* only after delay *t*Δ_*ij*_^1^, then produces a jump in current that decays exponentially with time constant *τ*_*s*_. The delay is a programmable parameter that increases the expressivity of a spiking neuron [8]. However, for the primary analysis using the covering-number, we treat this as a fixed parameter.

#### A.1.1 Spike Trains

Spike trains encode each input by converting its value into the timing and amplitude of Dirac delta pulses, which are then injected into the network to initiate the spiking process. The spike train is derived through a procedure that maps real-valued inputs to a tensor of Dirac delta functions.

Consider:

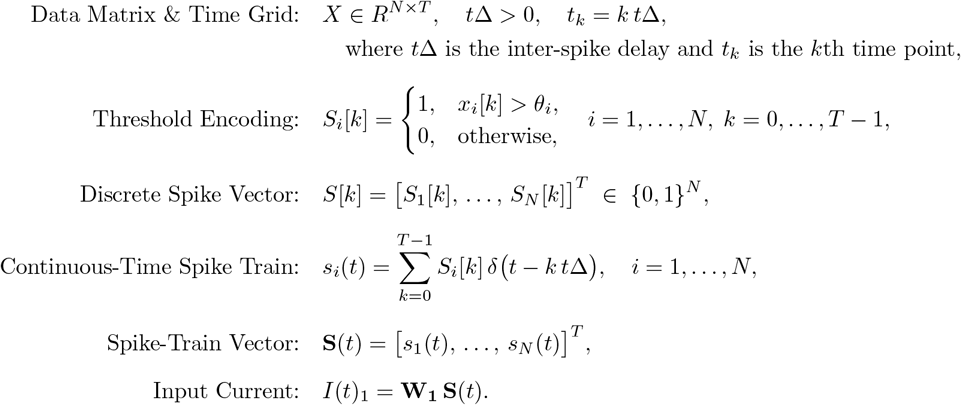

This process achieves two goals^2^. First, it binarizes the raw inputs into a sequence of 0*/*1 spikes. Second, it embeds those spikes in continuous time by replacing each “1” in the discrete tensor with a Dirac delta at the appropriate moment. Concretely, for neuron *i* we write

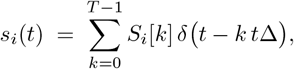

so that a “1” at index (*i, k*) becomes a δ-spike at time *t* = *k t*Δ. From the network’s point of view, this is equivalent to feeding it a stream of binary tensors *S*[0], *S*[1], …, *S*[*T* 1]. By casting those tensors as continuous spike trains, we naturally capture all temporal dependencies—so the LIF membrane and synaptic decay dynamics require no extra bookkeeping, because the δ pulses encode precisely when inputs arrive.

Once the membrane potential reaches the threshold *θ*, the neuron emits a spike and its potential is reset:

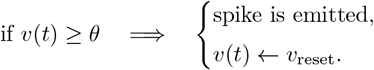

Importantly, the entire membrane potential process is initiated by the input state *spike trains*: at each time step *k*, the threshold-ed vector *S*[*k*] determines which neurons fire—every neuron with *S*_*i*_[*k*] = 1 emits a Dirac-δ spike, kick-starting the propagation of signals through the network.

#### A.1.2 Model Output

In order to actually use these networks it is important to understand how to retrieve output values from them. The output layer is monitored differently depending on the task being performed. For Example, during classification tasks, we monitor the output layer over a fixed time window *T*. Let

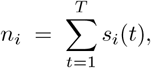

denote the spike count of output neuron *i* in the window *T*, where T represents the total elapsed time in the network. We map these rates to class probabilities with a softmax:

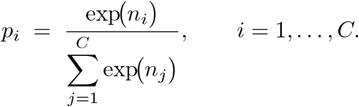

For sequence–generation tasks (e.g. language modeling) we take a snapshot of the output layer’s membrane potentials at the end of the time window. This moment is when the last token propagates entirely through the network.

Let

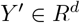

be this membrane potential vector and

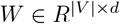

the embedding matrix whose *v*-th row represents vocabulary token *v*. The logit for token *v* is

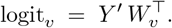

The probability assigned to token *i* is obtained with a soft-max:

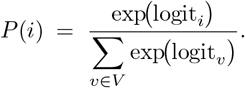

Once the spiking outputs are mapped to a probability distribution, training proceeds almost exactly as in a ANN.

#### A.1.2 Surrogate Gradient Descent

The commonly used Heaviside activation function is not differentiable, making it impossible to perform gradient descent. A commonly used method is to use a differentiable function which approximates the SNN activation function and use backpropagation through time (BPTT) to learn the weights of the network.

**Figure 1.**
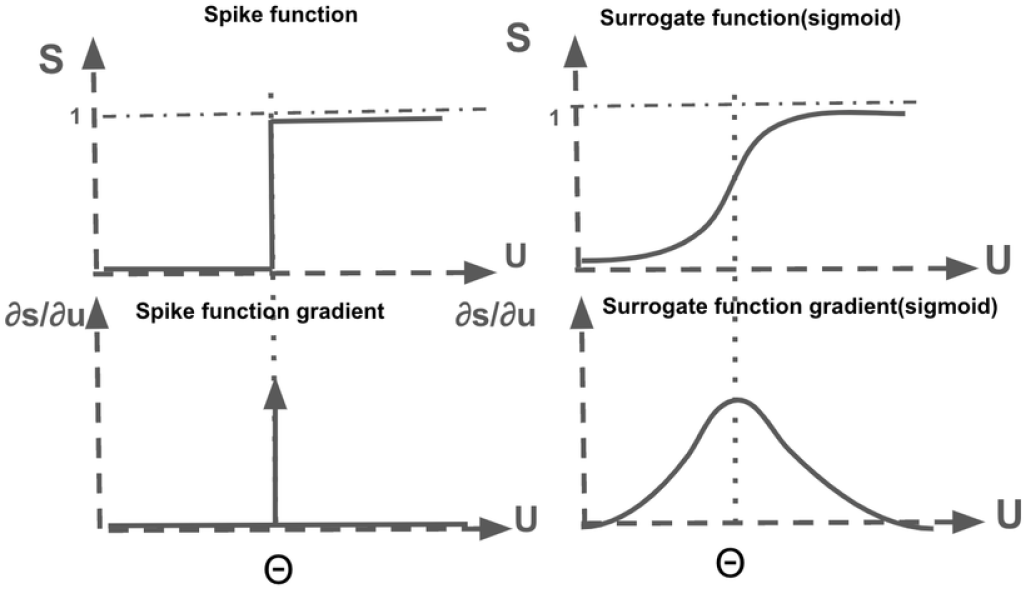
Visualization of Different Commonly Used Surrogate Gradients [20]

This allows the cross-entropy loss gradient to propagate through the network and update the weight values. Some networks also allow for learnable decay functions, but in the traditional LIF the decay rates and thresholds are fixed.

## B VC Dimension Proof Sketches for Supplementary Results of SNNs

We give informal sketches of the VC-dimension bounds of SNNs that help motivate our work. These sketches are not intended as full formal proofs (see the original references for complete arguments), but rather as a reader’s guide to the key ideas and constructions.

### Theorem B.1.

*For each n one can construct a feedforward spiking-neuron network (SNN) N with O*(*n*) *edges whose VC dimension is* Ω(*n*^2^). *This remains true even if all synaptic weights and firing thresholds are held fixed*.

*Proof Sketch*. Let

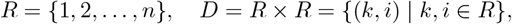

and fix any binary labeling stored in the *n* × *n* matrix *B* = [*b*_*k,i*_]. Encode each row *k* by the delay

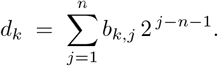

**Inputs**. The network has 2*n* + 1 input neurons: {*X*_1_, …, *X*_*n*_} (row selectors), {*Y*_1_, …, *Y*_*n*_} (column selectors), and *Z* (global reference). On presentation of (*e*_*k*_, *e*_*i*_), exactly *X*_*k*_, *Y*_*i*_, and *Z* fire at *t* = 0. The *X*_*k*_ spike then arrives at the first module *M*_*n*_ after delay *d*_*k*_.

### Module *M*_*m*_ (bit-extraction)

Each module *M*_*m*_ (*m* = *n, n* − 1, …, 1) does the following:

- *IN1:* data spike at

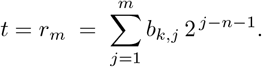
- *IN2:* reference spike from *Z* at *t* = 0.
- *OUT2:* if *b*_*k,m*_ = 1, emits at *t* = *r_m_*
- *OUT1:* emits at

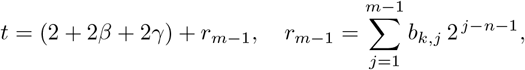

and forwards this to *M*_*m*−1_.

### Coincidence detection

For each column *i*, neuron *C*_*i*_ receives:

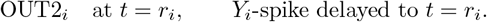

With unit weights and threshold 2, *C*_*i*_ fires exactly at *t* = *r*_*i*_ if and only if *b*_*k,i*_ = 1. No other *C*_*j*_ can fire.

Thus each (*k, i*) is correctly labeled by whether *C*_*i*_ spikes, and since we can choose all *n*^2^ bits arbitrarily via the delays {*d*_*k*_}, the network shatters *D* using only *O*(*n*) edges [11].

While these constructions are stated for non-leaky neurons, they rely solely on programmable delays and hence carry over to leaky membranes without loss of generality. Moreover, since adding programmable weights and thresholds cannot reduce expressivity, the same Ω(*n*^2^) lower bound holds for fully programmable SNNs. The authors also show that any SNN with *L* edges, programmable weights, delays, and thresholds, and depth *D* satisfies

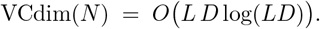

In the case of a network whose depth scales with its width (*D* = Θ(*L*)), this upper bound simplifies as follows:

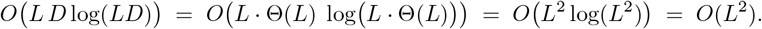

## C Causal Pieces & Local Lipschitz Continuity

This section provides an overview of the work done by Dold et al. Our work is focused on extending the implications of their work thus key points which were not defined in the main paper and the key local Lipschitz continuity proof sketch are in this section.

### Definition 2.

***Non Leaky Integrate-&-Fire (nLIF)*** *is a version of the LIF model in which there is no membrane leak. Specifically:*

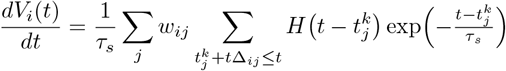

### Theorem C.1

(Local Lipschitz Continuity of nLIF Spike Times)

*Let N*_0_ ∈ *N and consider a single non-leaky integrate-and-fire (nLIF) neuron with input spike times*

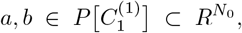

*where* 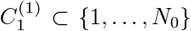 *is its causal set (i*.*e. the indices of all pre-synaptic spikes that occur before the output spike), synaptic time constant τ*_*s*_ > 0, *threshold* ϑ, *and weights satisfying*

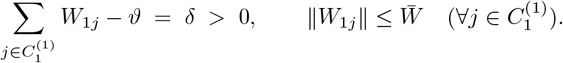

*Denote by*

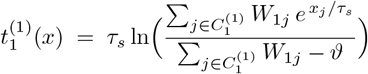

*the output spike time given inputs* 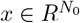. *Then on the causal piece* 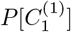,

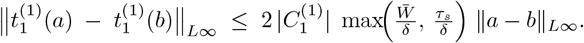

*Moreover, as long as one remains in a region where* ∑_*j* ∈*C*_*′ W*_1*j*_ −ϑ > 0, *continuity is preserved; if the causal set becomes empty the spike time can jump discontinuously*.

*Proof Sketch*.

*Within a causal piece, differentiability*. On 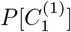, the causal set 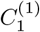 is fixed, so 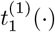 is a composition of logarithm, exponential, sums and quotients of affine functions of the inputs and weights. Hence it is smooth there.

One checks directly from

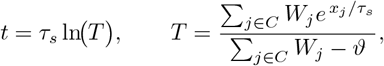

that for each input coordinate

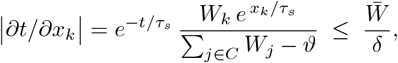

and for each weight

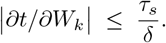

By the multivariate mean-value theorem in the *ℓ*_∞_–norm,

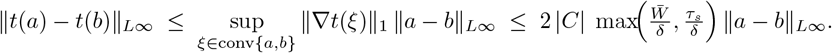

Here the factor 2 |*C*| arises since there are |*C*| partials in the inputs and |*C*| in the weights.

When the input or weights cross into a different causal set *C*^*′*^, continuity persists as long as ∑_*j* ∈ *C*_*′ W*_*j*_ −ϑ > 0. If one enters a region where no pre-synaptic input suffices to reach threshold, the output spike time jumps to ∞.

This completes the proof sketch of the work done by Dold et al. For a more formal overview of their work see the original paper [10]

## D Lipschitz Function Composition Example

### Example D.1.

Consider a vocabulary *M* where each *m*_*i*_ ∈ *M* is a word in an embedding space s.t. *m*_*i*_ is a *R*^*D*^ tensor. WLOG, these words are adjectives with positive or negative sentiment, such as (Happy, Sad, Good, Bad,…). The goal is to build a network that classifies these words based on their associated sentiment.

Using Time to First Spike (TTFS), a Lipschitz continuous spike encoding function can be created to encode the spike trains^1^. For simplicity a simple normalization TTFS function is chosen where

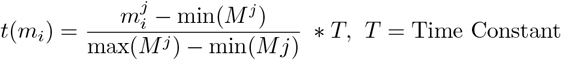

This encodes an input tensor where all input layer neurons will fire at some time *t*(*m*_*i*_)^*j*^. We assume that the *t*(*m*_*i*_)^*j*^ cells are close enough together s.t. that one input does not trigger a neuron to spike multiple times, so as not to invalidate our SIS setup. Clearly, this rate encoding scheme is Lipschitz continuous. Since the nLIF spike train inputs are bounded by 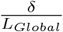 to ensure Lipschitz continuity with the *M* input space we construct a bound for our input samples s.t. 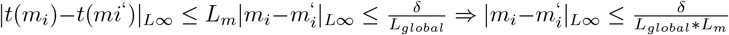 where *L*_*m*_ is the spike train encoding Lipschitz value and *L*_*global*_ is the global Lipschitz value for the nLIF w.r.t spike times and weights of the nLIF. The goal of this Example is to classify positive and negative sentiment from single input spikes. Let the output layer consist of two neurons, “pos” and “neg,” with spike times *T*_pos_, *T*_neg_. Define

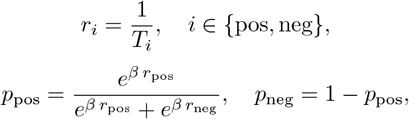

and classify by

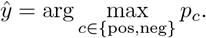

This decoding function converts these spike times back to outputs. This is an application of the softmax in the output and this decoding function is Lipschitz with some bound *L*_*o*_. Since the nLIF SNN is able to be decomposed into a set of Lipschitz continuous functions, the entire network is Lipschitz continuous s.t. for some *m*_*i*_, *m*_*j*_ ∈ *M* where

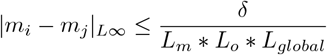

Then

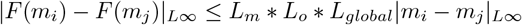

## Footnotes

1 The same pseudo-attention phenomena in an LIF exist in an nLIF. We focus on the LIF to avoid redundancy.

1 When we say RNN, we are referring to a vanilla RNN

2 pre-synaptic here simply means the immediate previous layer. This work uses a telescoping method to extend the Lipschitz bounds

1 A time delay represents the time for a signal to be transmitted from a pre-synaptic neuron to a post-synaptic neuron. These are constant throughout the network and mimic time to travel in actual neuronal communication

2 Threshold encoding is for illustrative purposes and is not Lipschitz continuous. Do not use this technique in the function composition in Section 3.

1 Not all spike encoding functions are Lipschitz continuous without added assumptions. Many like threshold encoding are not without added assumptions about the input space.

## References

[1] R.-J. Zhu, Q. Zhao, G. Li, and J. K. Eshraghian, “Spikegpt: Generative pre-trained language model with spiking neural networks,” arXiv preprint 2302.13939, 2023.

[2] M. Schmitt, “Vc dimension bounds for networks of spiking neurons.” in ESANN, 1999, pp. 429–434.

[3] P. Koiran and E. D. Sontag, “Vapnik-chervonenkis dimension of recurrent neural networks,” Discrete Applied Mathematics, vol. 86, no. 1, pp. 63–79, 1998.

[4] Y. Chen, H. Liu, K. Shi, M. Zhang, and H. Qu, “Spiking neural network with working memory can integrate and rectify spatiotemporal features,” Frontiers in Neuroscience, vol. 17, p. 1167134, 2023.

[5] A. Taherkhani, A. Belatreche, Y. Li, G. Cosma, L. P. Maguire, and T. M. McGinnity, “A review of learning in biologically plausible spiking neural networks,” Neural Networks, vol. 122, pp. 253–272, 2020.

[6] B. L. Edelman, S. Goel, S. Kakade, and C. Zhang, “Inductive biases and variable creation in self-attention mechanisms,” in International Conference on Machine Learning. PMLR, 2022, pp. 5793– 5831.

[7] M. Schmitt, “On the sample complexity of learning for networks of spiking neurons with nonlinear synaptic interactions,” IEEE Transactions on Neural Networks, vol. 15, no. 5, pp. 995–1001, 2004.

[8] W. Maass and M. Schmitt, “On the complexity of computing and learning with networks of spiking neurons,” in Electronic Proceedings of the Fifth International Symposium on Artificial Intelligence and Mathematics, http://rutcor.rutgers.edu/ãmai/. Journal version to appear in Information and Computation. Citeseer, 1998.

[9] A. M. Zador and B. A. Pearlmutter, “Vc dimension of an integrate-and-fire neuron model,” in Proceedings of the ninth annual conference on Computational learning theory, 1996, pp. 10–18.

[10] D. Dold and P. C. Petersen, “Causal pieces: analysing and improving spiking neural networks piece by piece,” 2025. [Online]. Available: https://arxiv.org/abs/2504.14015

[11] W. Maass and M. Schmitt, “On the complexity of learning for spiking neurons with temporal coding,” Information and Computation, vol. 153, no. 1, pp. 26–46, 1999.

[12] M. Chen, X. Li, and T. Zhao, “On generalization bounds of a family of recurrent neural networks,” arXiv preprint 1910.12947, 2019.

[13] W. Dorrell, M. Yuffa, and P. E. Latham, “Meta-learning the inductive bias of simple neural circuits,” in International Conference on Machine Learning. PMLR, 2023, pp. 8389–8402.

[14] A. L. Hodgkin and A. F. Huxley, “The dual effect of membrane potential on sodium conductance in the giant axon of loligo,” The Journal of physiology, vol. 116, no. 4, p. 497, 1952.

[15] A. L. Hodgkin and A. F. Huxley, “A quantitative description of membrane current and its application to conduction and excitation in nerve,” The Journal of physiology, vol. 117, no. 4, p. 500, 1952.

[16] D. J. Heeger, “Normalization of cell responses in cat striate cortex,” Visual neuroscience, vol. 9, no. 2, pp. 181–197, 1992.

[17] M. Carandini and D. J. Heeger, “Normalization as a canonical neural computation,” Nature reviews neuroscience, vol. 13, no. 1, pp. 51–62, 2012.

[18] P. D. King, J. Zylberberg, and M. R. DeWeese, “Inhibitory interneurons decorrelate excitatory cells to drive sparse code formation in a spiking model of v1,” Journal of Neuroscience, vol. 33, no. 13, pp. 5475–5485, 2013.

[19] X. Liu, “Inhibition snn: unveiling the efficacy of various lateral inhibition learning in image pattern recognition,” Discover Applied Sciences, vol. 6, no. 11, p. 611, 2024.

[20] E. O. Neftci, H. Mostafa, and F. Zenke, “Surrogate gradient learning in spiking neural networks: Bringing the power of gradient-based optimization to spiking neural networks,” IEEE Signal Processing Magazine, vol. 36, no. 6, pp. 51–63, 2019.

